# JADE family proteins regulate proteasome abundance and activity

**DOI:** 10.1101/2021.10.01.462752

**Authors:** LK Ebert, S Bargfrede, K Bohl, HT Cohen, R-U Müller, T Benzing, B Schermer

## Abstract

The JADE protein family (JADE1/2/3) has been implicated in a broad range of cellular functions, including WNT signaling, cell cycle control, regulated cell death, and transcriptional regulation through histone acetyltransferase complexes. All three paralogs share high sequence similarity and contain two PHD zinc finger domains. JADE1 has additionally been associated with cilia-related proteins and genetic disorders affecting kidney architecture and function. Despite their widespread expression, the molecular roles of JADE proteins remain incompletely understood.

Here, we identify JADE proteins as regulators of proteasome abundance and activity. Using kidney cells as a model, we demonstrate that loss of any single JADE protein led to a marked upregulation of almost all components of the 26S proteasome. Regulation occurred at the post-translational level and was not the consequence of increased transcription. Consistent with a role for JADE proteins in regulating overall proteasomal abundance, JADE-deficient cells displayed elevated proteasomal activity, while ectopic expression of JADE1, JADE2, or JADE3 reduced proteasomal function. Co-immunoprecipitation experiments confirmed the interaction between JADE1 and multiple proteasomal subunits, supporting a direct role of JADE proteins in modulating proteasome turnover, stability and abundance.

These findings reveal a novel function of JADE proteins in proteostasis and suggest that their previously reported cellular roles may, in part, be mediated through regulation of the ubiquitin-proteasome system.

## Introduction

The JADE protein family (JADE1/2/3) comprises zinc finger proteins containing plant homeodomains (PHDs) and is involved in diverse cellular processes. To date, however, most functional studies have focused on JADE1. JADE1 (PHF17; Gene for Apoptosis and Differentiation in Epithelia 1) was initially identified as an interactor of the tumor suppressor pVHL in a yeast two-hybrid screen (1). It exists in two isoforms: a long isoform (JADE1L) and a short isoform (JADE1S), which differ in their C-terminal regions due to splicing. JADE1S is the most extensively characterized isoform to date and has been the focus of most molecular studies (2). Mutations in *VHL* cause the autosomal dominant von-Hippel Lindau (VHL) disease, a hereditary tumor syndrome associated with hemangioblastomas, pheochromocytomas and clear-cell renal cell carcinoma (ccRCC). Aberrations of *VHL* are also found in the majority of sporadic ccRCCs (3–6). JADE1 has been shown to be stabilized by pVHL and this stabilization is diminished in the presence of pathogenic *VHL* mutations (1, 7–9). Moreover, JADE1 has been described as a single unit E3 ubiquitin ligase targeting β-catenin for degradation, thereby inhibiting canonical WNT signaling (10). Both elevated β-catenin levels and reduced JADE1 expression are associated with poor prognosis in RCC patients (7). This supports the current notion of JADE1 as a putative tumor suppressor in the kidney, related to pVHL.

In addition, JADE1 forms a protein complex with members of the NPH protein family, in particular NPHP4. Mutations in the *NPHP* genes result in Nephronophthisis (NPH), an autosomal recessive cystic kidney disease leading to kidney failure in children and adolescents (11). Similar to pVHL, NPHP4 stabilizes JADE1 and enhances inhibition of canonical WNT signaling (12). JADE1 has also been functionally linked to polycystin-1 (PC1) (13), encoded by the *PKD1* gene. Mutations in *PKD1* cause autosomal dominant polycystic kidney disease (ADPKD), the most frequent cystic kidney disease in adults. Since PC1, NPH proteins, and pVHL all localize to primary cilia, diseases caused by mutations in these genes are classified as ciliopathies—a group of disorders arising from ciliary dysfunction (14). Notably, JADE1 has been shown to co-localize with NPH proteins at the ciliary base (12).

Besides its role in ciliopathy-related signaling and WNT regulation, JADE1 has been described as a co-factor for two histone acetyltransferases (HATs), HBO1 and its homologue TIP60, thereby inducing histone H4 acetylation (15, 16). This implicates JADE1 in transcriptional and epigenetic regulation, DNA metabolism, and cell cycle regulation (15, 17, 18). Surprisingly, a gene trap mouse model targeting *Jade1* showed no major overt phenotype (19) which was recently confirmed in another *Jade1*-deficient mouse line (20). These observations suggest that additional proteins may functionally compensate for the loss of JADE1. The additional members of the Jade protein family, JADE2 (PHF15) and JADE3 (PHF16), are obvious candidates for this redundancy (19). The genes encoding JADE1, JADE2, and JADE3 are located on three different chromosomes (chr. 4, chr. 5, chr. X in the human genome and chr. 3, chr. 11 and chr. X in the mouse genome). As we demonstrate in detail, the three JADE proteins share a high sequence similarity and conserved protein domains (21).

Although little is known about specific functions of JADE2 and JADE3, emerging evidence suggests they may participate in related pathways: Like JADE1, JADE2 was proposed to act as an E3-ubiquitin ligase targeting the histone H3K4 demethylase LSD1 (22–24). JADE3 has been shown to be induced by WNT/β-catenin and may have oncogenic properties in colon cancer (25). More recently, JADE3 was found to regulate the antiviral gene interferon-induced transmembrane protein 3 (IFITM3), promoting NF-kB activation and restricting influenza A virus infection (26). Notably, all three JADE proteins are part of the HBO1 complex (15, 18, 27).

While JADE family members have been implicated in numerous pathways, their molecular functions have not been directly compared in the same tissue or cell type. Here, we investigate shared and distinct functions of *Jade1*, *Jade2*, and *Jade3* in differentiated murine renal tubular epithelial cells. Using CRISPR/Cas9-mediated genome editing, we generated monoclonal loss-of-function cell lines, employing two independent strategies per *Jade* gene to yield two deficient cell lines per gene. After confirming the loss of JADE expression and the absence of compensatory changes at the mRNA or protein level, we performed unbiased proteomic, transcriptomic and in-depth bioinformatic analyses. These revealed increased proteasome abundance and activity as a common molecular phenotype of JADE-deficient cells.

### Experimental Procedures

#### Plasmids and antibodies

Plasmids containing *JADE1S*, *VHL*, and *EPS*^1-225^ cDNA were previously described (12, 28). *JADE1L*, *JADE2* and *JADE3* cDNA were obtained by PCR from *Human* Embryonic Kidney (HEK) 293T cDNA. Murine *Jade1S*, *Jade1L*, *Jade2*, *Jade3*, *Psmb4*, and *Psmd2* cDNA were obtained by PCR from murine inner-medullary collecting duct (mIMCD3) cDNA. cDNA fragments were inserted into a modified pcDNA6 vector (Invitrogen) containing an N-terminal FLAG-tag using standard cloning techniques. For the generation of the CRISPR/Cas9-mediated mutant cell lines the selected single guide (sg)RNAs were cloned into pSpCas9(BB)-2A-GFP (PX458), gifted from Feng Zhang (Addgene plasmid #48138) (29). All plasmids were verified by Sanger sequencing.

Primary antibodies were obtained from Sigma-Aldrich (monoclonal mouse anti-FLAG (M2), F3165, 1:10000; polyclonal rabbit anti-FLAG, F7425, 1:2000) and Serotec (monoclonal mouse anti-V5, MCA1360, 1:5000). Secondary antibodies were obtained from Jackson ImmunoResearch (goat anti-rabbit HRP-coupled, 111-035-003; goat anti-mouse HRP-coupled, 115-035-003; both 1:30000).

#### Cell culture and transfections

mIMCD3 (ATCC CRL-2123) cells were cultured in DMEM-F12 (Sigma-Aldrich) supplemented with 10% fetal bovine serum (FBS) and 2 mM GlutaMAX (Gibco). For FACS sorted cell lines the medium was additionally supplemented with 100 U/ml Penicillin-Streptomycin (Gibco). mIMCD3 subclone 8 (sc.#8) was selected after FACS sorting the parental line into single cells for a higher genetic homogeneity. For the generation of the CRISPR/CAS9-mediated mIMCD3 sc.#8 mutant cell lines, the cells were seeded in 6-well plates, grown to 50 % confluency and transfected with Lipofectamine 2000 (Invitrogen) according to the manufacturer’s instructions. For the siRNA transfection mIMCD3 sc.#8 cells were grown to 60% confluency. The cells were transfected with Lipofectamine RNAiMAX according to the manufacturer’s instructions. The final concentration of each siRNA (Dharmacon, L-059773-01-0005/L-059777-01-0005/L-065325-01-0005) was 10 nM. HEK293T cells were cultured in DMEM (Gibco) supplemented with 10% FBS. For transfection experiments in HEK293T cells, the cells were seeded in 10 cm dishes and grown to 50-60% confluency. Plasmid DNA was transiently transfected using the calcium phosphate method. 24 h after transfection cells were harvested in 8 ml of cold PBS, centrifuged and the cell pellet was used for lysis and subsequent coimmunoprecipitation. All cells were cultured at 37°C in the presence of 5% CO_2_ and tested negative for mycoplasmas.

#### Coimmunoprecipitation

HEK293T cell pellets were resuspended in 1 ml of lysis buffer (20 mM Tris (pH 7.5), 1% (v/v) TritonX-100, 50 mM NaCl, 15 mM Na_4_P_2_O_7_, 50 mM NaF) and incubated for 15 min at 4 °C. After homogenization the cell suspension was centrifuged (20,000 x g, 15 min, 4 °C) and then the supernatant was ultracentrifuged (100,000 x g, 30 min, 4 °C). 50 µl of the supernatant after ultracentrifugation was used as lysate (input) and boiled with 2x Laemmli buffer. Anti-FLAG agarose beads (M2, Sigma) were added to the remaining supernatant. After incubation for 1 h at 4 °C the beads were extensively washed with lysis buffer and boiled in Laemmli buffer. Proteins were resolved by SDS-Page.

#### Generation of mutant cell lines

For each member of the Jade family two sgRNAs were designed using the CRISPR finder tool (http://www.sanger.ac.uk/htgt/wge/)(30) and Benchling (http://benchling.com). The sgRNA sequences are shown in supplemental table S1. Cloning of the sgRNA containing plasmids and transfection are described above. 48 h after transfection GFP-positive cells were sorted in 96-well plates on a BD FACSAria^TM^ III. After expanding the colonies arising from single cells, the first screening for mutants was performed using PCR, followed by Sanger sequencing. Primer sequences can be obtained from supplemental table S1. Validation was done by qPCR and a parallel reaction monitoring (PRM) assay.

#### qPCR

mIMCD3 wild type, Jade mutant cells, and siRNA transfected cells were washed with PBS and RNA extraction was performed with the Direct-zol RNA Miniprep kit (Zymo Research) following the manufacturer’s instructions including a DNase1 treatment step. This was followed by reverse transcription using the High-Capacity cDNA Reverse Transcription kit (Applied Biosystems). *Jade1/2/3* mRNA was assessed by SYBR Green (Thermo Fisher Scientific) qPCR using *Hprt1* as endogenous control. Primers are listed in supplemental table S1. The qPCR experiments were performed on a QuantStudio 12K Flex Real_time PCR System (Thermo Fisher Scientific).

#### PRM assays

JADE protein expression has been difficult to detect by shotgun proteomics experiments. Therefore, we decided to use a PRM assay (31). Preparation of the samples for targeted mass spectrometry was performed as previously described (32). All samples were measured by the proteomics facility at CECAD. All samples were analyzed on an Orbitrap Exploris 480 mass spectrometer coupled to an EASY nLC 1200 UPLC (both Thermo Scientific). Peptides were loaded with solvent A (0.1% formic acid in water) onto an in-house packed analytical column (30 cm × 75 μm I.D., filled with 2.7 μm Poroshell EC120 C18 (Agilent)). Peptides were chromatographically separated at a constant flow rate of 300 nl min^-1^ using with the following gradient: initial 4 % solvent B (0.1% formic acid in 80% acetonitrile), ramp to 25% B within 72 min, to 55% B within 8 min and to 95% B within 2 min, followed by washing and column equilibration. The Exploris was equipped with a FAIMS Pro Interface (Thermo) set to a CV of −47 and operated in scheduled PRM mode. The MS1 survey scan was acquired from 400 to 1000 m/z at a resolution of 120,000. Target masses were isolated in a 2 h window and fragmented by HCD with collision energy of 30%. The AGC target was set to 100 % and resulting spectra recorded with a resolution of 120,000. Product ions were detected in the Orbitrap at a resolution of 17,500.

Target list generation and follow-up analysis were performed in Skyline 20.2. Unique target peptides and their retention times were chosen based on their detectability in data-independent trials using the identical setup and gradient. Results were compared against their theoretical spectra simulated by the Prosit algorithm (33) plugin in Skyline (34, 35). Only areas with a dotp value above 0.7 were included in the analysis to ensure the confidence of the result. The targets are listed in supplemental table S2.

##### Proteome analyses

For each replicate, a 10 cm dish of mIMCD3 cells of the indicated genotype was harvested and snap-frozen. Pellets were resuspended in Urea buffer (8 m Urea, 50 mm ammoniumbicarbonate, Halts Protease-phosphatase-inhibitor (Thermo Scientific)). After clearing of the sample (16000g, 30 min at 4°C) the lysates were reduced (10 mM Dithiothreitol, 1 h, RT) and alkylated (50 mM Chloroacetamide, 30 min, RT, in the dark). An equal amount of protein was diluted to reach a concentration of 2M urea and subjected to tryptic digestion (enzyme:substrate ratio of 1:50). On the next day, double-layered stage-tip clean up (C18) was performed.

All samples were analyzed on a Q Exactive Plus Orbitrap (Thermo Scientific) mass spectrometer that was coupled to an EASY nLC (Thermo Scientific). Peptides were loaded with solvent A (0.1% formic acid in water) onto an in-house packed analytical column (50 cm — 75 µm I.D., filled with 2.7 µm Poroshell EC120 C18, Agilent). Peptides were chromatographically separated at a constant flow rate of 250 nL/min using the following gradient: 3-4% solvent B (0.1% formic acid in 80 % acetonitrile) within 1.0 min, 4-27% solvent B within 119.0 min, 27-50% solvent B within 19.0 min, 50-95% solvent B within 1.0 min, followed by washing and column equilibration. The mass spectrometer was operated in data-dependent acquisition mode. The MS1 survey scan was acquired from 300-1750 m/z at a resolution of 70,000. The top 10 most abundant peptides were isolated within a 1.8 Th window and subjected to HCD fragmentation at a normalized collision energy of 27%. The AGC target was set to 5e5 charges, allowing a maximum injection time of 55 ms. Product ions were detected in the Orbitrap at a resolution of 17,500. Precursors were dynamically excluded for 30.0 s.

All mass spectrometric raw data were processed with Maxquant (version 1.5.3.8) using default parameters. Briefly, MS2 spectra were searched against the canonical Uniprot mouse fasta database (UP000000589; May 4, 2020) and the MaxQuant default list of common contaminants. False discovery rates on protein and PSM level were estimated by the target-decoy approach to 1% (Protein FDR) and 1% (PSM FDR) respectively. The minimal peptide length was set to 7 amino acids and carbamidomethylation at cysteine residues was considered as a fixed modification. Oxidation (M) and Acetyl (Protein N-term) were included as variable modifications. The match-between runs option was enabled. Student’s T-tests were calculated in Perseus (version 1.6.1.1) after removal of decoys and potential contaminants. Data were filtered for at least 3 out of 3 values in at least one condition. Remaining missing values were imputed with random values from the left end of the intensity distribution using Perseus defaults.

##### Interactome analysis

For each replicate, a 10 cm dish of *Jade1* KO1 mIMCD3 cells was transfected with 10 µg of either FLAG-tagged *Jade1S*, *Jade1L* or an empty vector control using Lipofectamine (Invitrogen) 2000 following the manufacturer’s instructions. 48 h after transfection cells were harvested and lyzed (50 mM Tris (pH 7.5), 150 mM NaCl, 0.5% (w/v) Sodium deoxycholate, 1% (v/v) Triton X-100) for 15 min at 4°C. After sonication (Bioruptor, 10 min, cycle 30/30 sec) and centrifugation (4°C, 15 min) anti-FLAG agarose beads (M2, Sigma) were added to the supernatant and incubated at 4°C overnight. After extensive washing of the beads, 5% SDS in 1xPBS was added and proteins were released from the beads by incubation at 95°C for 3 min. Subsequently, the supernatant was reduced with DTT and alkylated with CAA followed by two single-pot solid-phase-enhanced sample preparations (SP3).

All samples were analyzed on a Q Exactive Plus Orbitrap (Thermo Scientific) mass spectrometer that was coupled to an EASY nLC (Thermo Scientific). Peptides were loaded with solvent A (0.1% formic acid in water) onto an in-house packed analytical column (50 cm — 75 µm I.D., filled with 2.7 µm Poroshell EC120 C18, Agilent). Peptides were chromatographically separated at a constant flow rate of 250 nL/min using the following gradient: 3-5% solvent B (0.1% formic acid in 80 % acetonitrile) within 1.0 min, 5-30% solvent B within 65.0 min, 30-50% solvent B within 13.0 min, 50-95% solvent B within 1.0 min, followed by washing and column equilibration. The mass spectrometer was operated in data-dependent acquisition mode. The MS1 survey scan was acquired from 300-1750 m/z at a resolution of 70,000. The top 10 most abundant peptides were isolated within a 1.8 Th window and subjected to HCD fragmentation at a normalized collision energy of 27%. The AGC target was set to 5e5 charges, allowing a maximum injection time of 110 ms. Product ions were detected in the Orbitrap at a resolution of 35,000. Precursors were dynamically excluded for 15.0 s.

All mass spectrometric raw data were processed with Maxquant (version 1.5.3.8) using default parameters. Briefly, MS2 spectra were searched against the canonical Uniprot MOUSE_UP000000589.fasta (downloaded at: 26.08.2020) database, including a list of common contaminants. False discovery rates on protein and PSM level were estimated by the target-decoy approach to 1% (Protein FDR) and 1% (PSM FDR) respectively. The minimal peptide length was set to 7 amino acids and carbamidomethylation at cysteine residues was considered as a fixed modification. Oxidation (M) and Acetyl (Protein N-term) were included as variable modifications. The match-between runs option was enabled for replicates from the same group. LFQ quantification was enabled using default settings. Student’s t-tests were calculated in Perseus (version 1.6.1.1) after removal of decoys and potential contaminants. Data were filtered for at least 4 values in at least one condition.

#### In vitro assay of 26S proteasome activity

The proteasomal activity assay was performed as previously described (36). For each replicate with the mIMCD3 cells, the cells were grown into 6-well plates and grown until 70% confluency. HEK29T cells were grown into 6-well plates, transfected at 50% confluency with 2 µg of DNA using the calcium phosphate method, and harvested 24 h after transfection. Cells were collected in proteasomal activity assay buffer (50 mM Tris-HCL, pH 7.5, 10% glycerol, 5 mM MgCl_2_, 0.5 mM EDTA, 2 mM ATP, 1 mM DTT) and immediately frozen in liquid nitrogen. Before performing the assay, samples were lysed and centrifuged at 10000 g for 10 min at 4°C. 25 µg of protein was loaded on a flat-bottom, black fluorescence 96-well microplate and incubated with the fluorogenic substrate (Z-Gly-Gly-Leu-AMC, BML-ZW8505-0005, ENZO Life Science). Fluorescence (360 nm excitation, 430 nm emission) was monitored every 5 min for 2 h at 37°C using a fluorescent plate reader (EnSpire Multimode Plate Reader, Perkin Elmer).

#### mRNA sequencing

Total RNA extraction was performed with the direct-zol RNA Miniprep kit (Zymo Research) following the manufacturer’s instructions. Library preparation and sequencing was performed by the Cologne Center for Genomics. Libraries were prepared using the Illumina® TruSeq® mRNA stranded sample preparation Kit. Library preparation started with 300 ng total RNA. After poly-A selection, mRNA was purified and fragmented using divalent cations under elevated temperature. The RNA fragments underwent reverse transcription using random primers, followed by second strand cDNA synthesis with DNA Polymerase I and RNase H. After end repair and A-tailing, indexing adapters were ligated. The products were then purified and amplified to create the final cDNA libraries. After library validation and quantification (TapeStation 4200, Agilent), equimolar amounts of library were pooled and quantified by using the Peqlab KAPA Library Quantification Kit and the Applied Biosystems 7900HT Sequence Detection System. The pool was sequenced on an Illumina NovaSeq6000 with PE100 read length and a minimum of 35 million reads per sample.

The reads were trimmed with Trimmomatic version 0.36 (37) using default parameters. The trimmed reads were mapped to the GRCm39 mouse reference genome with STAR version 2.6 (38) using default parameters. mRNA expression was analyzed using DESeq2 (version 1.28.1) (39) package of R (version 4.0.0) software (https://www.R-project.org/). A p-value cut-off of <0.05 was applied to the list of differentially regulated mRNAs after pairwise comparison of the *Jade*-deficient cell lines with wild type. P-values were adjusted for multiple testing.

#### Experimental design and statistical rationale

To ensure data reproducibility, we used 3 biological replicates for the proteome study, the PRM assay as well as the transcriptome study. For the interactome we used 4-5 biological replicates. Detailed information about filtering and statistical analysis are under the respective subsections. For the proteasome activity assay samples were assayed in technical triplicates and in biological triplicates. Statistical significance was calculated with a one-way ANOVA with Dunnett’s post-hoc test (* p ≤ 0.05, ****p ≤ 0.0001). For the qPCR analysis an unpaired Student’s 2-tailed t-test was used to calculate statistical significance by using GraphPad Prism version 10 (GraphPad Software, San Diego, CA). Statistical significance is represented as follows: *P < 0.05, **P < 0.01, and ***P < 0.001. The results displayed are an average of at least 3 biological replicates, and error bars indicate standard error of mean (SEM).

## Results

### JADE protein family members share common interactors related to renal disease

JADE proteins share two PHD zinc finger domains, multiple PEST domains, and nuclear localization signals (Fig. 1A). Based on these structural similarities, we investigated whether JADE2 and JADE3 share molecular functions with JADE1. JADE1 has been characterized as an interactor of the ciliary protein pVHL (8) and of the nephrocystin protein complex (12). Therefore, we cloned the cDNA of human JADE2 and JADE3 in expression vectors and performed coimmunoprecipitation experiments, which included the short protein isoform JADE1S as positive control. V5-tagged pVHL (V5.VHL) was co-expressed together with JADE1S (F.Jade1S), JADE2, JADE3, or a control protein, all with an N-terminal Flag-tagged. These experiments revealed pVHL to co-precipitate with all three JADE proteins (Fig. 1B). We performed similar experiments with V5.NPHP4 (Fig. 1C) and V5.NPHP1 (Fig. 1D). Again, both NPHP4 and NPHP1 specifically co-precipitated with all the JADE proteins. Additionally, coimmunoprecipitation experiments show that FLAG-tagged JADE1S and JADE2 are each able to interact with a V5 tagged version of the same protein (Fig. 1E,F) and are capable to homodimerize. Based on the high structural similarity we checked for heterodimerization. Coimmunoprecipitation of JADE2 with JADE1 or JADE3 suggested that these proteins may exist in common protein complexes (Fig. 1E). In summary, all JADE proteins associate with relevant renal protein complexes, very similar to their putative common role as co-factors and components of the HBO1/HAT complex (15, 27), indicating that they could interact in the same complex with a common function.

**Figure 1:**
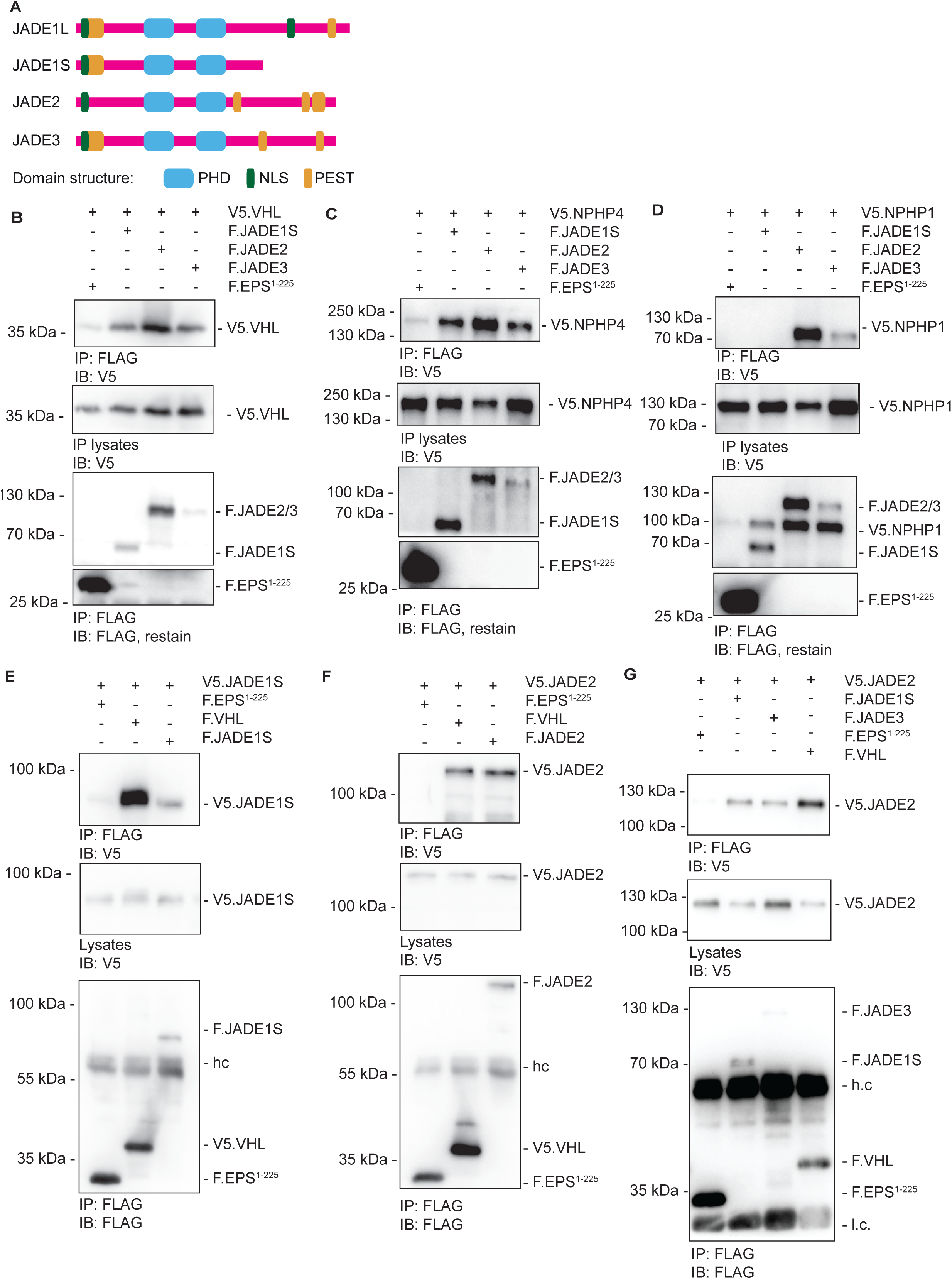
All JADE protein family members co-precipitate with the ciliopathy proteins NPHP1, NPHP4, and pVHL. **(A)** Domain structure of the human JADE proteins. PHD – Cys4HisCys3 plant homeo domain; NLS – nuclear localization signal; PEST – protein degradation amino acid sequence. Domain information obtained from EMBOSS/epestfind (57), SMART(58), and NLS prediction (consensus sequence K – (K/R) –X – (K/R)). **(B-D)** HEK293T cells were transiently transfected with **(B)** V5-tagged pVHL (V5.VHL), **(C)** V5-tagged Nphp4 (V5.NPHP4), and **(D)** V5-tagged Nphp1 (V5.NHPH1), and either FLAG-tagged JADE1S (F.JADE1S), Jade2 (F.JADE2), Jade3 (F.JADE3) or a control protein (F.EPS^1-225^). Immunoprecipitates with an anti-FLAG antibody were analyzed by immunoblotting with antibodies (anti-V5; anti-FLAG) as indicated. V5.VHL, V5.NPHP1, and V5.NPHP4 all coimmunoprecipitate with F.Jade1S/2/3 but not with F.EPS^1-225^. Blots are representative for at least three independent experiments. **(E,F)** HEK293T cells were transiently transfected with **(E)** V5.JADE1S and **(F)** and V5.JADE2 and either FLAG-tagged JADE1S/2, pVHL, or EPS^1-225^. Immunoprecipitates with an anti-FLAG antibody were analyzed by immunoblotting with antibodies (anti-V5; anti-FLAG) as indicated. V5.Jade1S/2 both coimmunoprecipitate with F.JADE1S/2 and F.VHL but not with F.EPS^1-225^. **(G)** HEK293T cells were transiently transfected with V5.JADE2 and either FLAG-tagged JADE1S, JADE3, pVHL, or EPS^1-225^. Immunoprecipitates with an anti-FLAG antibody were analyzed by immunoblotting with antibodies (anti-V5; anti-FLAG) as indicated. V5.JADE2 coimmunoprecipitate with F.JADE1S, F.JADE3, and F.VHL but not with F.EPS^1-225^ (h.c.: heavy chain, l.c.: light chain).

### JADE family members show ubiquitous transcription with spikes in different tissues

The N-terminal part of the JADE proteins shows the highest level of similarity as illustrated by the higher protein identity score between JADE2/3 and JADE1S compared to JADE1L (Supplemental Fig. S1A) (21). We used available data sets of the Human Protein Atlas to analyze tissue expression (40). Protein expression data was only available for JADE1 and shows global expression in most tissues with squamous epithelial cells of the oral mucosa, esophagus, and uterine, as well as hepatocytes and kidney tubules expressing the highest levels (Supplemental Fig. S1B). mRNA expression data support the ubiquitous expression of *JADE1* in most tissues with higher levels in the pancreas and endocrine and female tissues (Fig. S1C). *JADE2* appears ubiquitously expressed in most tissues with enrichment in the brain, endocrine tissues, female tissues, and lymphoid tissue. The overall *JADE3* expression is relatively low as compared to *JADE1* and *JADE2*. The expression pattern is similar to *JADE2* with the highest levels in the brain, endocrine tissues, female and male tissues, and lymphoid tissues (Supplemental Fig. S1C). Importantly, while the expression of all three JADE genes is confirmed in most tissues, there is generally one JADE family member predominantly expressed (e.g., JADE1 in pancreas, bone marrow, or breast; JADE2 in cerebral cortex, lymph nodes, and midbrain; JADE3 in epididymis, placenta, and testis). In addition, the *JADE* family is conserved across species and has a target identity matching the mouse orthologue to the human sequences of 92.45 %, 91.44 %, and 81.92 %, for JADE1, JADE2, and JADE3, respectively (41). Single-cell transcriptomic data from mouse kidneys revealed that all Jade proteins are expressed in the kidney with a predominance of *Jade1* over *Jade2* and *Jade3* and with co-expression of all three proteins in cells of the collecting duct (Supplemental Fig. S1D) (42). Therefore, we decided to target *Jade* expression in this specific cell type.

### Neither inactivation of a single JADE gene nor combined downregulation of two family members resulted in compensatory expression changes in the others

To analyze and compare transcriptional and translational consequences of Jade protein deficiency, we used the established mIMCD3 (murine inner medulla collecting duct 3) cell line and targeted individual *Jade* genes with CRISPR/CAS9. We designed two individual non-overlapping sgRNAs (single-guide RNAs) for each of the *Jade* genes to account for potential off-target effects. The first sgRNA was targeted to the region encoding the first PHD domain, the second sgRNA to the downstream exon containing the second PHD domain’s start (Supplemental Fig. S2A). Mutant monoclonal cell lines were screened and validated by PCR and Sanger sequencing; the resulting mutant Jade protein structures are depicted in Supplemental Fig. S2B. When performing qPCR analyses for the different *Jade* genes, we observed a significant reduction of mRNA expression of the targeted Jade gene in the respective mutant cell line (Fig. 2A-C). Targeted proteomic PRM assays confirmed the absence of individual JADE protein expression in the specific Jade-deficient cell lines (Fig. 2D-F). Remarkably, we did not observe any compensatory upregulation of the remaining Jade family members on transcript and protein level. To address whether a compensatory regulation is dependent on the lack of two out of three of the Jade family members, we transfected mIMCD3 cells with siRNA targeted against *Jade1*, *Jade1* and *Jade2*, *Jade1* and *Jade3*, or *Jade2* and *Jade3* and evaluated transcript levels with qPCR. Similar to the results in the *Jade*-deficient cell lines we also did not observe any compensation upon loss of two of the *Jade* family members (Fig 2G).

**Figure 2:**
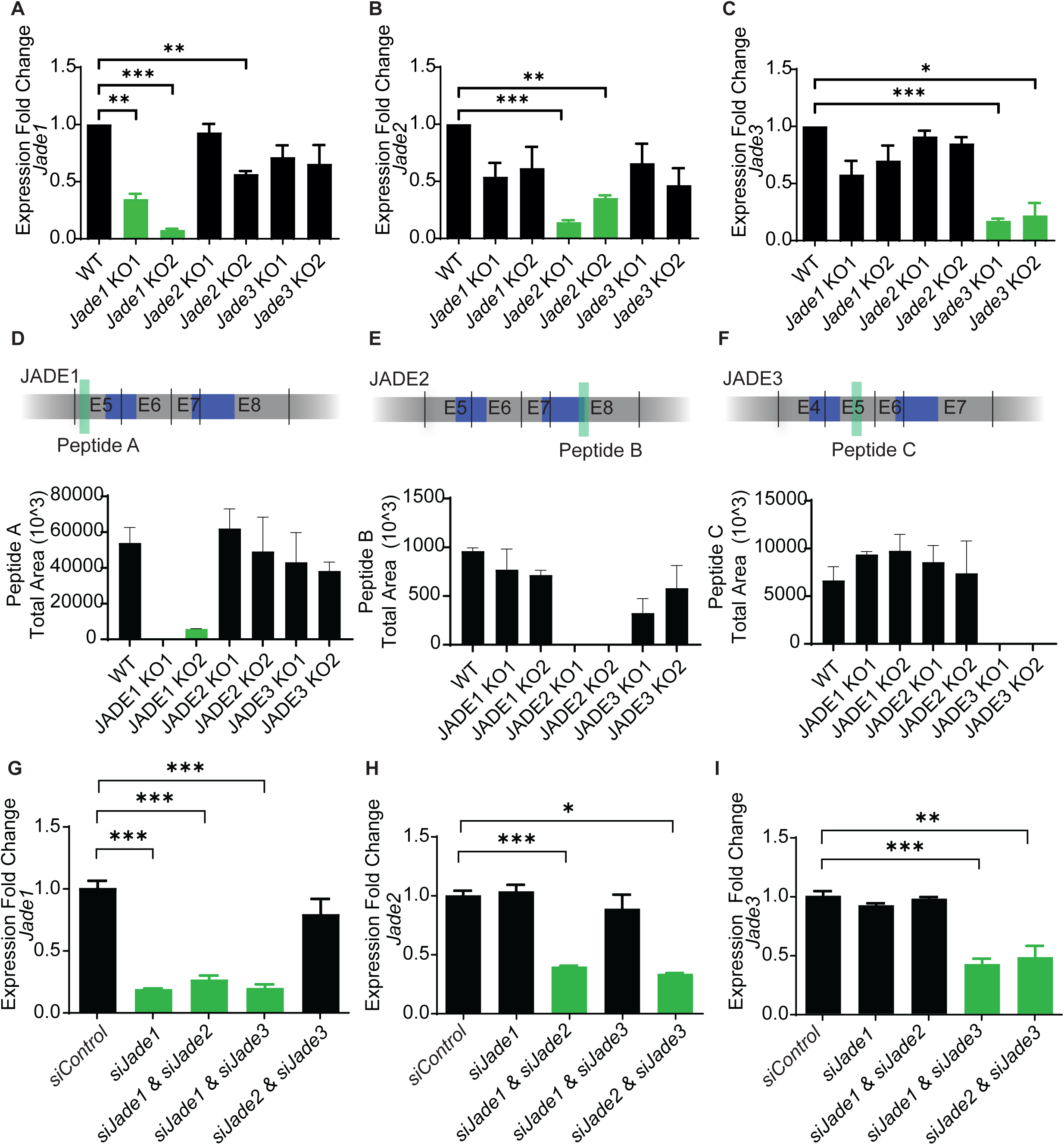
Deletion of one specific Jade protein affects neither transcription nor expression of the remaining family members. **(A-C)** qPCR data comparing the expression fold change of each targeted gene across the different cell lines. Only the expression of the targeted gene in the respective mutant cell lines is significantly reduced, while the other Jade mRNAs are not significantly altered. (N=3, students t-test, * p ≤ 0.05, ** p ≤ 0.01, *** p ≤ 0.001) **(D-F)** Parallel reaction monitoring (PRM) assay for **(D)** Jade1, **(E)** Jade2, and **(F)** Jade3 comparing the peak areas for the selected peptides across mIMCD3 WT and Jade mutant cell lines for protein quantification. The localization of peptides selected for PRM assay is depicted above the bar charts. (N=3). (G-I) mIMCD3 were treated with the indicated siRNA combinations for 48h. qPCR data comparing the expression fold change of each targeted gene across the different knock-down cell lines. Only the expression of the targeted gene of the respective siRNA shows a significant reduction, while the other Jade mRNAs are not significantly altered. (N=3, students t-test, * p ≤ 0.05, ** p ≤ 0.01, *** p ≤ 0.001)

### Proteomic analyses of JADE-deficient cell lines reveal an increased abundance of almost all proteasomal components

We next performed unbiased proteomic analysis from all six JADE-deficient cell lines and the wildtype control cells (Supplemental table S3). For each JADE-deficient cell line the principal component analysis showed the clustering of the replicates and the separation of WT versus the pair of mutant cell lines by the first principal component (Fig. 3A-C). When plotting the Student’s T-test difference of KO1 versus KO2 for each of the JADE family members we found that they correlate and that expression of most deregulated proteins was shifted in the same direction within both mutant cell lines for each JADE protein (Supplemental Fig. S3A-C). Taking the sum of the significantly regulated proteins for both mutant cells for each JADE protein, we then calculated the overlap of significantly regulated proteins between the three JADE paralogs (Fig. 3D). This revealed 644 proteins with significantly altered expression in all JADE1/2/3-deficient cell lines, supporting the concept that JADE paralogs share biological functions. A Gene Ontology and KEGG pathway analysis of these 644 proteins revealed terms related to the 26S proteasome, ribosomes, and protein translation to be enriched (Fig. 3E). Indeed, many proteins related to the proteasome were significantly upregulated in the cell lines, as shown in a representative scatter plot of the JADE1 KO2 cell line (Fig. 3F) and in a heat-map summarizing the results from all cell lines (Fig. 3G). We also observed an altered expression of ribosomal proteins, tRNA ligases, as well as translation initiation and elongation factors in a similar manner (Supplemental Fig. S3D-F), being suggestive of a high protein turnover.

**Figure 3:**
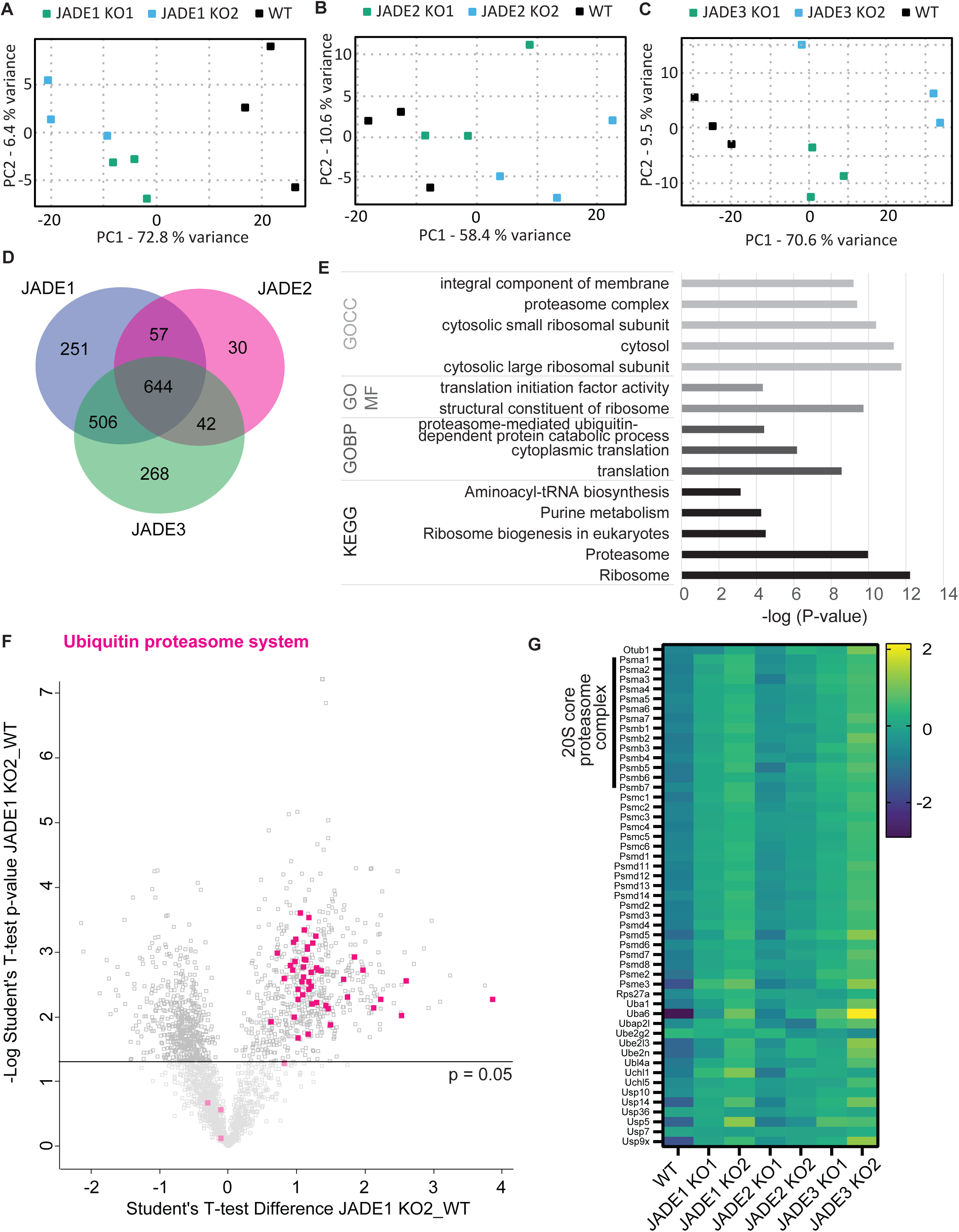
Proteomic analysis reveals upregulation of the proteasome in cell lines lacking any one of the Jade family members. (A-C) Principal component analysis (PCA) plot of the protein expression data of WT, Jade1 (A), Jade2 (B), and Jade3 (C) KO1 and KO2 cell lines. Depicted are the first two principal components (PC1 and PC2). The axes represent the percentages of variation explained by the principal components. The replicates of the different groups cluster together and the difference between mutant and wildtype groups explains the largest variance. (D) Venn diagram depicting the overlap between significantly regulated proteins across the Jade-deficient cell lines. The Jade1 section includes all proteins that are significantly regulated (p-value < 0.05) in either one or both of the Jade1-deficient cell lines. The Jade2 and Jade3 sections generated in the same way. (E) GO and KEGG pathway annotation based on a Fisher exact test of the 644 proteins found to be regulated in all Jade mutant cell lines. (F) Representative scatter plot with the t-test differences in protein expression of the Jade1 KO2 mutant cells vs the wildtype cells on the x-axis and the statistical significance (-log_10_ Student’s t-test p-value) on the y-axis. Proteins associated with the ubiquitin proteasome system are highlighted in magenta. (G) Heatmap of the proteins highlighted in (F) based on logarithmized LFQ values for all Jade-deficient cell lines and the control line visualizing the upregulation of proteasome-associated proteins.

### Elevated proteasomal activity in JADE deficiency

To investigate whether these changes in protein expression caused a higher proteasomal activity, we monitored the degradation of a fluorogenic substrate for the chymotrypsin-like activity of the proteasome. Indeed, all JADE-deficient cell lines showed a significant increase in proteasomal activity as compared to the wildtype cells (Fig. 4 A). Consistently, using exogenously expressed JADE proteins, we could show that the proteasome activity was reduced compared to an overexpressed control protein (Fig. 4B).

**Figure 4:**
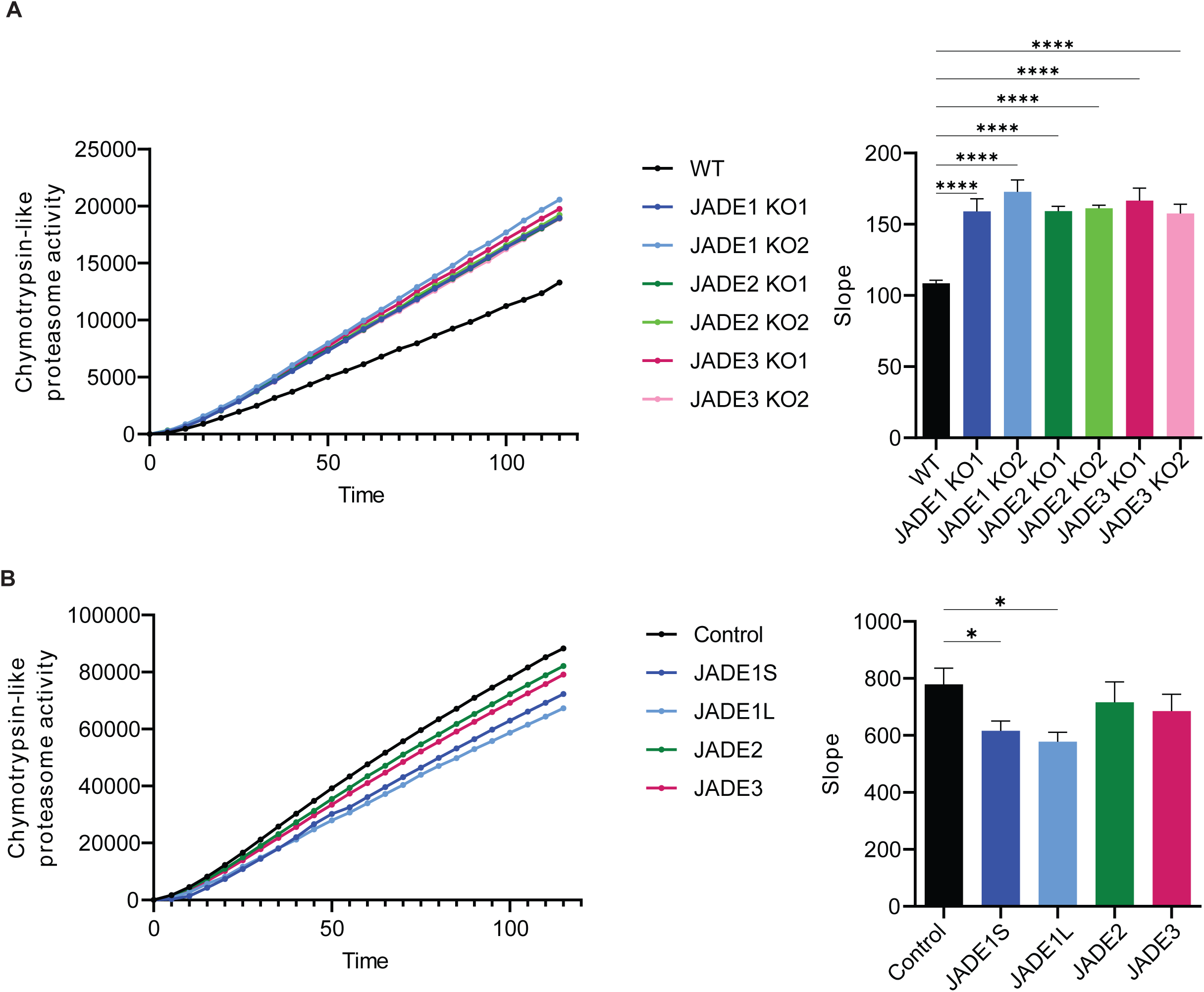
Increased proteasome activity observed in Jade-deficient cell lines. **(A)** Measurement of the proteasome activity in wildtype and Jade-deficient mIMCD3 cell lines. The proteasome activity of Jade1, Jade2, or Jade3-deficient cell lines is increased compared to the wildtype control. Quantification of the slope as readout for proteasome activity (N=3, one-way ANOVA with Dunnett’s post-hoc test, ****p ≤ 0.0001) **(B)** Measurement of the proteasome activity in HEK293T cell with either Jade1S, Jade1L, Jade2, Jade3, or Podocin (control) overexpressed. All cell lines with overexpressed Jade proteins show a decreased activity compared to the control with only Jade1S/L reaching statistical significance. Quantification of the slope as readout for proteasome activity (N=3, one-way ANOVA Dunnett’s post-hoc test, * p ≤ 0.05)

### Increased proteasome abundance is not caused by transcriptional regulation

Based on the role of *Jade1/2/3* as transcriptional and putative epigenetic regulators, we aimed to investigate whether the increased proteasome activity was caused by transcriptional upregulation of its components. To answer this question, we performed unbiased transcriptomic analyses with mRNA-Sequencing data from all six JADE-deficient cell lines and the control cell line (Supplemental table S4). For *Jade1*-deficient cell lines, principal component analysis showed a narrow clustering of the replicates and a clear separation of WT versus mutant cell lines accounting for 75 % of the variance (Supplemental Fig. S4A). For *Jade2* and *Jade3* the replicates clustered clearly together, the separation from the WT cell lines was based primarily on the second principal component (Supplemental Fig. S4B,C). When plotting the fold change for KO1 versus KO2 for *Jade1* we saw a clear correlation (Supplemental Fig. S4D). The same visualization for *Jade2* and *Jade3* revealed that the individual mutant cell lines for each of them are less correlated than seen for *Jade1* (Supplemental Fig. S4E,F). Taking the sum of the significantly regulated genes for each *Jade* family member, we found that 567 genes were commonly regulated within the family (Supplemental Fig. S5A). However, when we individually checked the expression of the proteasomal components across all cell lines we did not observe significant expression changes (Supplemental Fig. S5B,C). Additionally, we did not observe an upregulation in ribosomal genes and genes encoding for tRNA ligases or transcription initiation and elongation factors (Supplemental Fig. S5D-F). Taken together, the increased abundance and activity of the proteasome was not based on changes in RNA levels in JADE-deficient cells.

### Proteasomal core components co-precipitate with JADE1L and JADE1S

Searching for a direct connection between the 26S proteasome and Jade proteins, we re-expressed either FLAG-tagged JADE1S or JADE1L or a control protein in a JADE1-deficient cell line and performed an anti-FLAG coimmunoprecipitation experiment. Pulled-down protein complexes were analyzed by nLC-MS/MS (Supplemental table S5). The heatmap of iBAQs of the proteins that are exclusively found in the JADE1L/S samples shows JADE1 as the most abundant among these proteins (Fig. 5A). Plotting the Student’s T-test Difference between JADE1L and the control versus the –log Student’s T-test p-value, we observed not only several known JADE1 interactors (green) but also several components of the proteasome (magenta) significantly enriched in the JADE1L sample (Fig. 5B). We could confirm the same for JADE1S. Here, specifically PSMA1 was among the most enriched proteins (Fig 5C). Furthermore, we could show that JADE2 and JADE3 co-immunoprecipitate with PSMB4 and PSMD2 (Supplemental Fig. S6A,B). While these results indicate that the JADE proteins and components of the proteasome are part of the same complex, PSMB4 was previously identified as a strong interactor in a yeast-to-hybrid screen in which a truncation of JADE1 lacking both PHDs was used as bait (10) strengthening a direct connection between these proteins.

**Figure 5:**
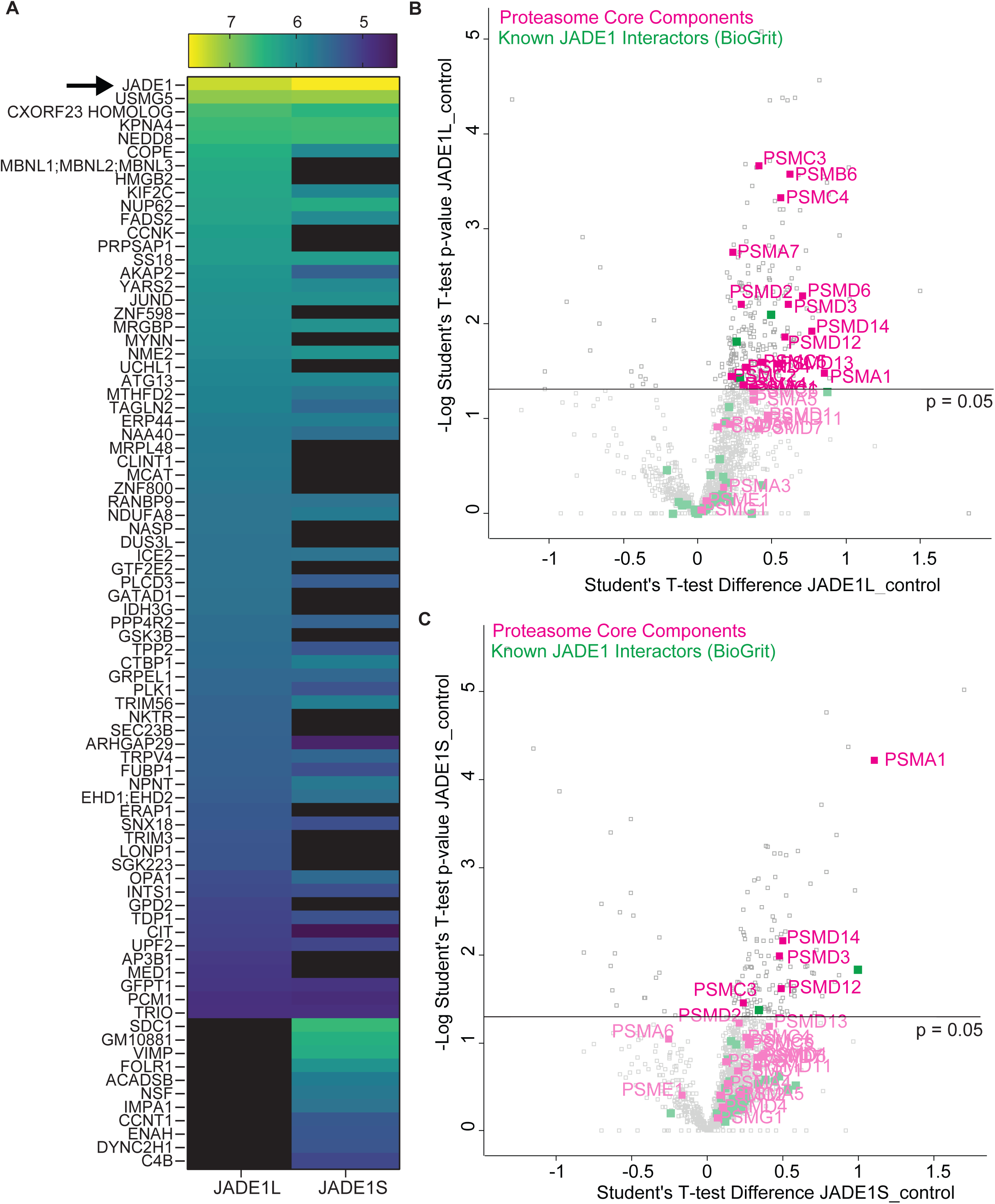
Interactome analysis reveals a link between Jade1L/S proteins and proteasomal components. **(A)** Jade1 KO1 mIMCD3 cells were transiently transfected with either Flag-tagged Jade1L, Jade1S, or a control plasmid. Pulldown was performed with anti-Flag (M2) sepharose beads. Heatmap of average iBAQ values for proteins which were never measured in the control samples, but in either all replicates for Jade1L, all replicated for Jade1S, or all replicates of both. The protein with the highest iBAQ is Jade1 for both, the Jade1S and the Jade1L pulldown. **(B/C)** Quantitative analysis of the Jade1L **(B)** & Jade1S **(C)** interactome. Proteins on the right side, above the p=0.05 line, are determined significantly enriched. Proteasomal components are highlighted in magenta, known JADE1/Jade1 interactors based on known interactors annotated in BioGrid (59) in green

## Discussion

JADE1 and, in part, the other JADE paralogs have been previously described as components of several protein complexes, and their potential cell biological functions have been mainly deduced by JADE interacting or associated proteins. Thus, JADE proteins were found to be part of the HBO1/HAT complex (15), to bind and become stabilized by pVHL, to directly ubiquitinate β-catenin and by that regulate canonical WNT signaling on multiple levels (8, 10, 12), or to be involved in cell cycle regulation through a PLK1/CK1 kinase module(43, 44). In addition, altered expression of JADE proteins has been identified in several tumor entities, in line with their protein-protein interactions with several tumor suppressors, including pVHL and ING4 (1, 7, 27). Until now, research on Jade proteins was primarily focused on the short isoform of JADE1 (JADE1S) (21) and not a single study has compared all three individual JADE proteins in the very same cellular context. In this study, we present for the first time an unbiased comparison of the proteomic and transcriptomic changes in JADE1, JADE2, and JADE3 deficient cell lines. Multiple previous studies suggested a specific role of JADE1 in renal ciliopathies (VHL, NPH (7, 8, 12)), ADPKD (13), renal cell carcinoma (7, 45, 46) and kidney regeneration after injury (47). Strikingly, we could confirm some of the key protein-protein interactions (pVHL, NPHP1 and NPHP4) related to renal ciliopathies for all three JADE proteins (Fig. 1). Therefore, we continued our work with the well-characterized renal tubular epithelial cell line (mIMCD3) to generate the JADE-deficient cell lines for the current comparative study. Importantly, previously published datasets confirmed that renal epithelial cells of the collecting duct express all JADE proteins *in vivo* (Supplemental Fig. S1D).

With CRISPR/CAS9 mediated genome editing, we generated two independent JADE-deficient cell lines for each JADE family member. These cell lines were validated by Sanger sequencing, qPCR analyses of mRNA expression, and a targeted proteomics approach (Fig. 2). Proteomic expression profiling revealed upregulated expression of more than 80% of the individual subunits of the 26S proteasome (Fig. 3G), and we could demonstrate increased proteasome activity accordingly, underlining the functional relevance of the altered proteasome abundance. This suggests that JADE proteins have a common function in balancing or tuning proteasome abundance. Since this is not caused by transcriptional regulation of proteasome expression as reflected by our RNA-Sequencing analyses (Supplemental Fig. S4), we conclude that JADE proteins might either modulate posttranslational modifications of individual proteasome components or, nonspecifically, induce the process of proteophagy. At this point, the finding of proteasomal subunits co-precipitating with JADE1S and JADE1L strongly suggests a direct involvement of JADE proteins in the targeting of the proteasome for degradation. While the detailed underlying mechanism of this regulation still needs elucidation, the observed high proteasomal activity in JADE deficient cells has several implications in the context of renal diseases. Enhanced proteasomal activity might increase the sensitivity of cells towards therapeutic inhibition of the proteasome. In Jade deficient cells, proteasome inhibition might more efficiently increase the unfolded protein burden and proteotoxic stress to toxic levels ultimately leading to apoptotic cell death. This could be relevant both for cancer cells with low or no JADE expression as well as in renal ciliopathies. Here, the disease genes pVHL, NPHP’s and Polycystin-1 have been demonstrated to stabilize JADE1 expression (1, 13) with mutant proteins lacking this specific effect (8, 13). In consequence, the low levels of JADE in mutant cells could cause increased proteasomal activity. Remarkably, the use of the proteasome inhibitor Carfilzomib has been shown to ameliorate the cystic kidney phenotype in a mouse model of ADPLD (48) by increasing cell death and decreasing proliferation of renal epithelial cells. In this model, Jade deregulation and the general high proteasomal activity might be one of the mechanisms targeted by Carfilzomib. Proteasome inhibitors have also been very recently identified among other drugs in a study screening about 8000 compounds for their effect on viability of PKD deficient cells (49). Further preclinical studies in mouse models are required to investigate under which exact conditions targeting the proteasome in cystic kidneys could be a useful option.

JADE1 is a protein localized at the ciliary transition zone and interacting with ciliopathy proteins, suggesting that the high abundance and activity of the proteasome might affect primary cilia. In all JADE-deficient cell lines, we do not observe any significant change in the number of ciliated cells (data not shown). The UPS, however, has been shown to be essential for the regulation of primary cilia. Inhibition of the proteasome with MG132 or depletion of proteasomal subunits with RNAi affects both ciliogenesis and ciliary disassembly (50). Interestingly, many proteasomal subunits have been identified as part of protein complexes pulled-down with transition zone and basal body proteins NPHP2, NPHP5 and NPHP8 (51) and with BBS proteins (52). Increased global proteasomal activity has not been directly linked to any ciliary function. One could speculate that increased proteasomal activity due to reduced JADE expression might positively affect primary cilia in renal ciliopathies. Interestingly, loss of cilia has been found to suppress cyst growth in ADPKD (53). As such, loss of cilia may indeed be positive in some renal ciliopathies and might be counteracted under certain conditions through depletion of JADE and subsequent upregulation of proteasomal activity.

## Abbreviations

ADPKD: autosomal dominant polycystic kidney disease
ADPLD: autosomal dominant polycystic liver disease
ccRCC: clear-cell renal cell carcinoma
FBS: fetal bovine serum
HAT: histone acetyltransferase
HEK293T: *Human* Embryonic Kidney 293T
iBAQ: intensity-based absolute quantification
JADE: gene for apoptosis and differentiation in epithelia
JADE1S: short protein isoform of JADE1
JADE1L: long protein isoform of JADE1
LFQ: label-free quantitation
mIMCD3: murine inner medullary collecting duct 3
NLS: nuclear localization signal
NPH: nephronophthisis
PC1: polycystin-1
PEST: protein degradation amino acid sequence
PHD: Cys4HisCys3 plant homeo domain
PRM: parallel reaction monitoring
SP3: single-pot solid-phase-enhanced sample preparation
UPS: ubiquitin-proteasome-system
VHL: von-Hippel Lindau

## Author contributions

L.K.B and B.S. conceptualization; L.K.E, H.T.C, R.-U.M methodology; L.K.E. and K.B. formal analysis; L.K.E. and S.B. investigation; L.K.E. visualization; L.K.E. and B.S. writing – original draft; L.K.E., K.B., H.T.C., R.-U.M, T.B., and B.S. writing – review and editing; B.S. ad R.-U.M. funding acquisition. All authors approved the final version of the paper.

## Acknowledgements

We thank Stefanie Keller and Martyna Brütting for excellent technical assistance and members of the laboratories for helpful discussions. In particular, we want to thank David Vilchez for his critical advice. We acknowledge the help of the CECAD proteomics core facility and the Cologne Center for Genomics, Cologne, Germany. This work was supported by the large instrument grant INST 1856/71-1 FUGG by the German Research Foundation (DFG Großgeräteantrag). We thank the FACS & IMAGING Core Facility at Max Planck Institute for Biology of Ageing, Cologne, for assisting with sorting cells for generation of Jade KO cell lines. This study was supported by the German Research Foundation (SCHE1562/8-1 to BS) and funded by the Deutsche Forschungsgemeinschaft (DFG, German Research Foundation) under Germany’s Excellence Strategy - EXC 2030 - 390661388.

## Data availability

The proteome and interactome mass spectrometry proteomics data have been deposited to the ProteomeXchange Consortium (http://proteomecentral.proteomexchange.org) via the PRIDE partner repository (54) and via Panorama Public (55). RNA sequencing data have been deposited in the ArrayExpress database at EMBL-EBI (www.ebi.ac.uk/arrayexpress) (56). Login information is available upon request.

## This article contains supplemental data

Supplemental figure S1-6

Supplemental table S1-5

