## Supplementary material for "JADE family proteins regulate proteasome abundance and activity": Suppl. Figures S1-S6 and Suppl. Tables S1-S2

**Supplemental material included:**

Supplemental Figure S1

Supplemental Figure S2

Supplemental Figure S3

Supplemental Figure S4

Supplemental Figure S5

Supplemental Figure S6

Supplemental table S1

Supplemental table S2

Supplemental table S3

Supplemental table S4

Supplemental table S5

### Supplemental figures

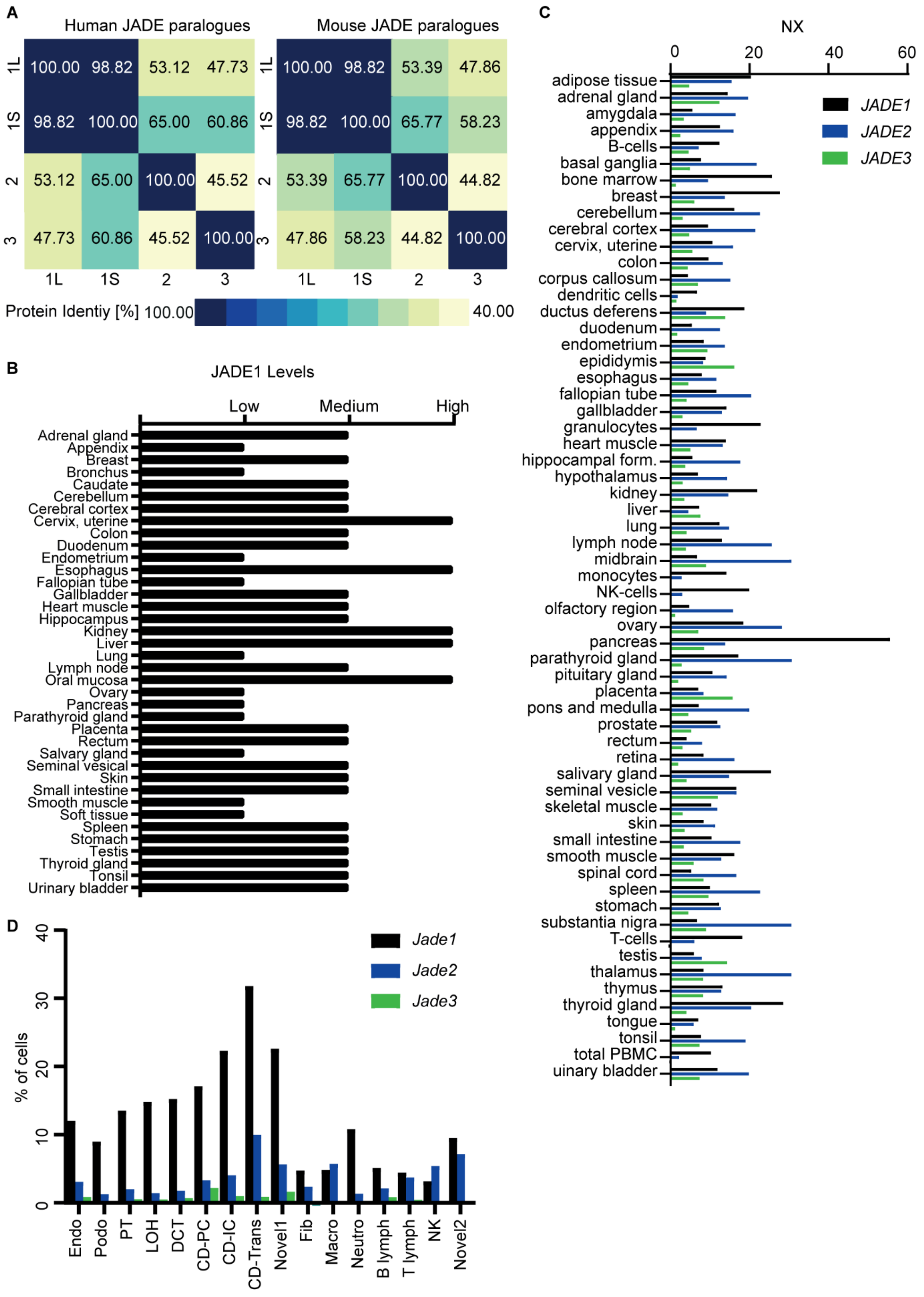

**Supplemental Figure S1: Sequence alignment and expression profiles of the JADE protein family members.** (A) Protein identity matrix for human and mouse JADE family paralogues. Protein sequences were obtained from Uniprot (1) and the identity score was calculated using Clustal Omega (2). (B) Protein Expression Levels for JADE1 across different tissues. Based on tissue micro arrays obtained from The Human Protein Atlas version 19.3 and Ensembl version 92.38. Proteomic data available from https://v19.proteinatlas.org/download/normal_tissue.tsv.zip (3). (C) RNA expression profiles across different tissues for *JADE1/2/3*. Consensus transcript expression levels based on transcriptomic data from HPA, GTEx and FANTOM5. Normalized expression (NX) based on the maximum NX for each gene in the different data sets. Data is obtained from The Human Protein Atlas v 19.3 and Ensembl version 92.38. Transcriptomic data available from https://v19.proteinatlas.org/download/rna_tissue_consensus.tsv.zip (3). (D) Gene expression data of single cell transcriptomics of mouse kidney. Shown is the percentage of cells expressing each *Jade* gene in the different cell clusters: endothelial, vascular, and descending loop of Henle (Endo), podocyte (Podo), proximal tubule (PT), ascending loop of Henle (LOH), distal convoluted tubule (DCT), collecting duct principal cell (CD-PC), collecting duct intercalated cell (CD-IC), collecting duct transitional cell (CD-Trans), fibroblast (Fib), macrophage (Macro), neutrophil (Neutro), lymphocyte (lymph), natural killer cell (NK)(4).

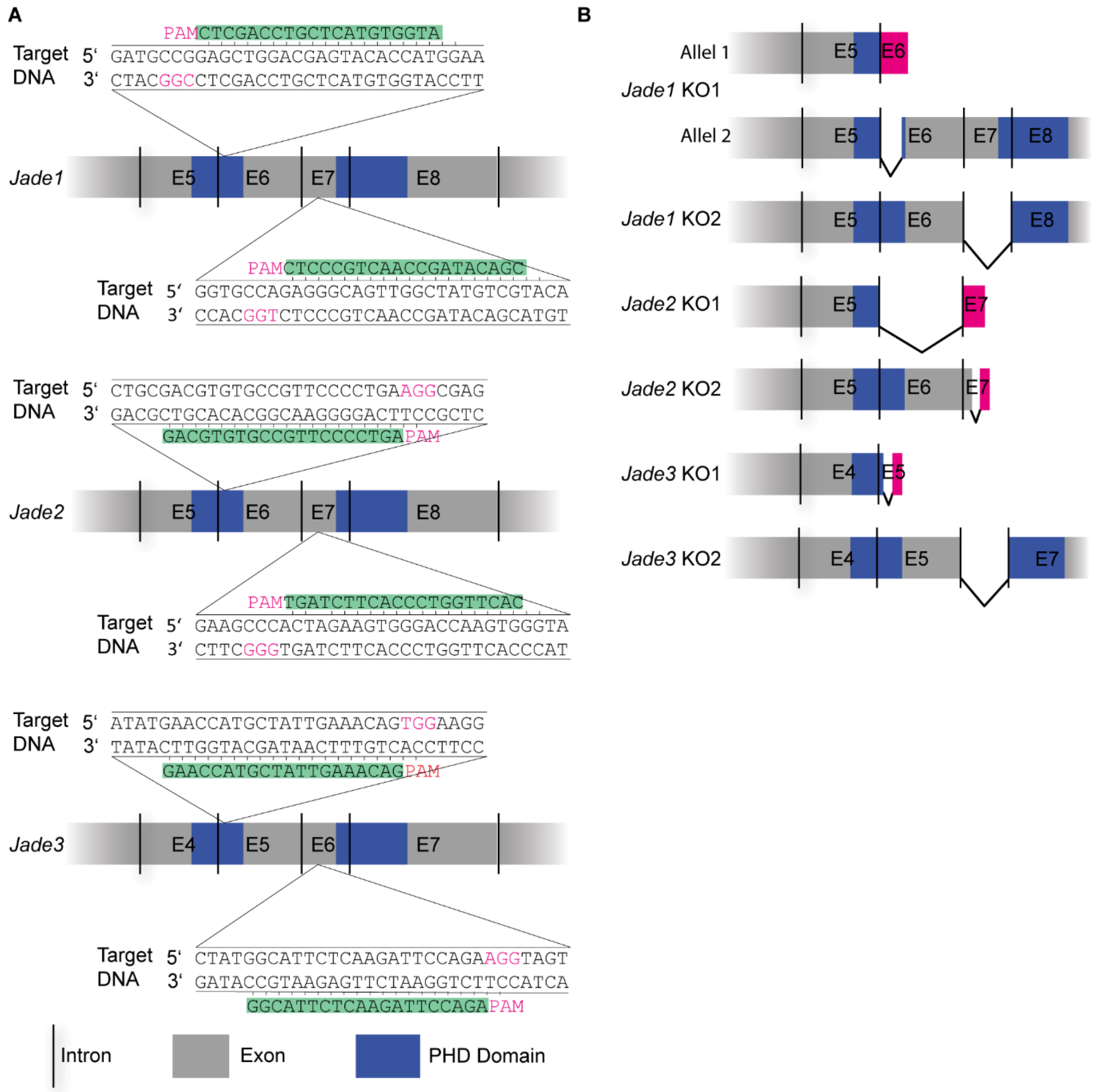


**Supplemental figure S2: Generation and validation of *Jade1/2/3* mIMCD3 mutant cell lines.** (A) Selection of sgRNAs targeting exons containing the first and the second PHD domain. sgRNA selection was based on location and off-target scores using Wellcome Sanger Institute Genome Editing (https://wge.stemcell.sanger.ac.uk/) (5). (B) Sanger sequencing of mRNA confirms mutations.

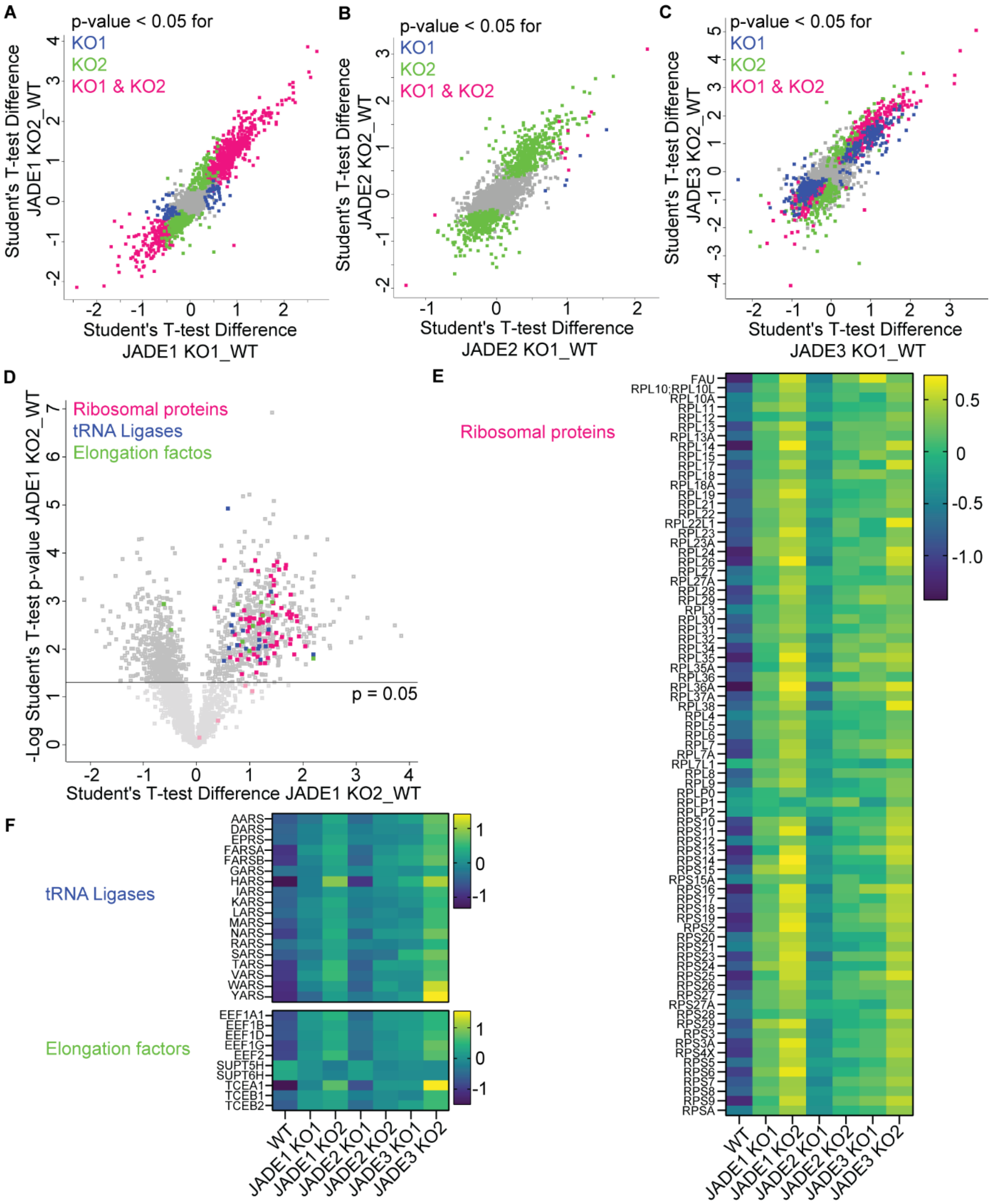


**Supplemental figure S3: Protein turnover machinery is upregulated upon loss of JADE family members. (A-C)** Comparison of the two different mutant cell lines for JADE1 **(A)**, JADE2 **(B)**, and JADE3 **(C)**. The fold change of KO1 is depicted on the x-axis, the fold change of the KO2 on the y-axis. Proteins are highlighted according to their significance as indicated in the figure. **(D)** Scatter plot with the t-test differences in protein expression of the JADE1 KO2 mutant cells vs the wildtype cells on the x-axis and the statistical significance (-$\log_{10}$ Student's t-test p-value) on the y-axis. Ribosomal proteins are highlighted in magenta, tRNA ligases in blue, and elongation factors in green. **(E, F)** Heatmap of the proteins highlighted in (D) based on logarithmic LFQ values for all seven cell lines to visualize the upregulation of ribosomal proteins (E), tRNA ligases, and elongation factors (F) across JADE-deficient cell lines.

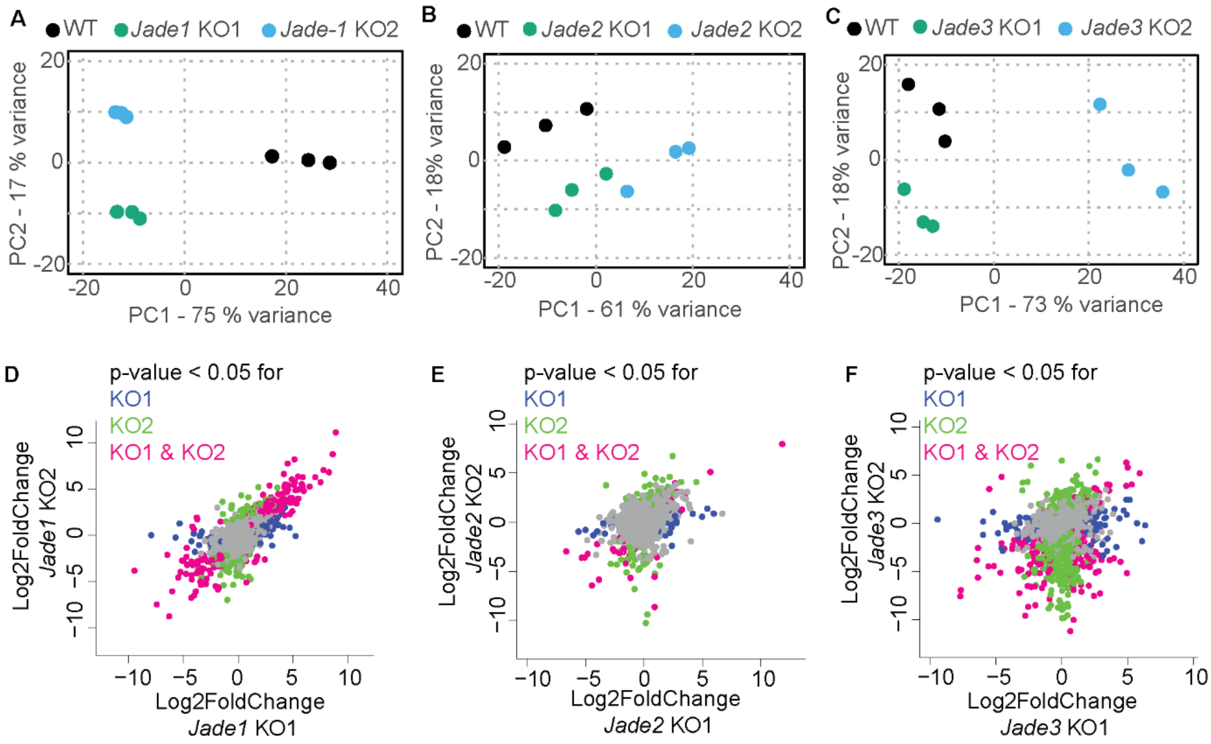


**Supplemental figure S4: Transcriptomic analysis reveals no major regulation of the proteasome. (A)** PCA plot of the gene expression data of WT, *Jade1* KO1 and KO2 cell lines. Depicted are the first two principal components (PC1 and PC2). The axes represent the percentages of variation explained by the principal components. The replicates of the different groups cluster together and the difference between mutant and wildtype groups explains the largest variance. PCA plots of *Jade2* **(B)** and *Jade3* **(C)** cell lines show clustering of the groups. Here, the second component accounts for the difference between mutant and wildtype cell lines. **(D-F)** Comparison of the two different mutant cell lines for *Jade1* **(D)**, *Jade2* **(E)**, and *Jade3* **(F)**. The fold change of KO1 is depicted on the x-axis, the fold change of the KO2 on the y-axis. Genes are highlighted according to their significance as indicated in the figure.

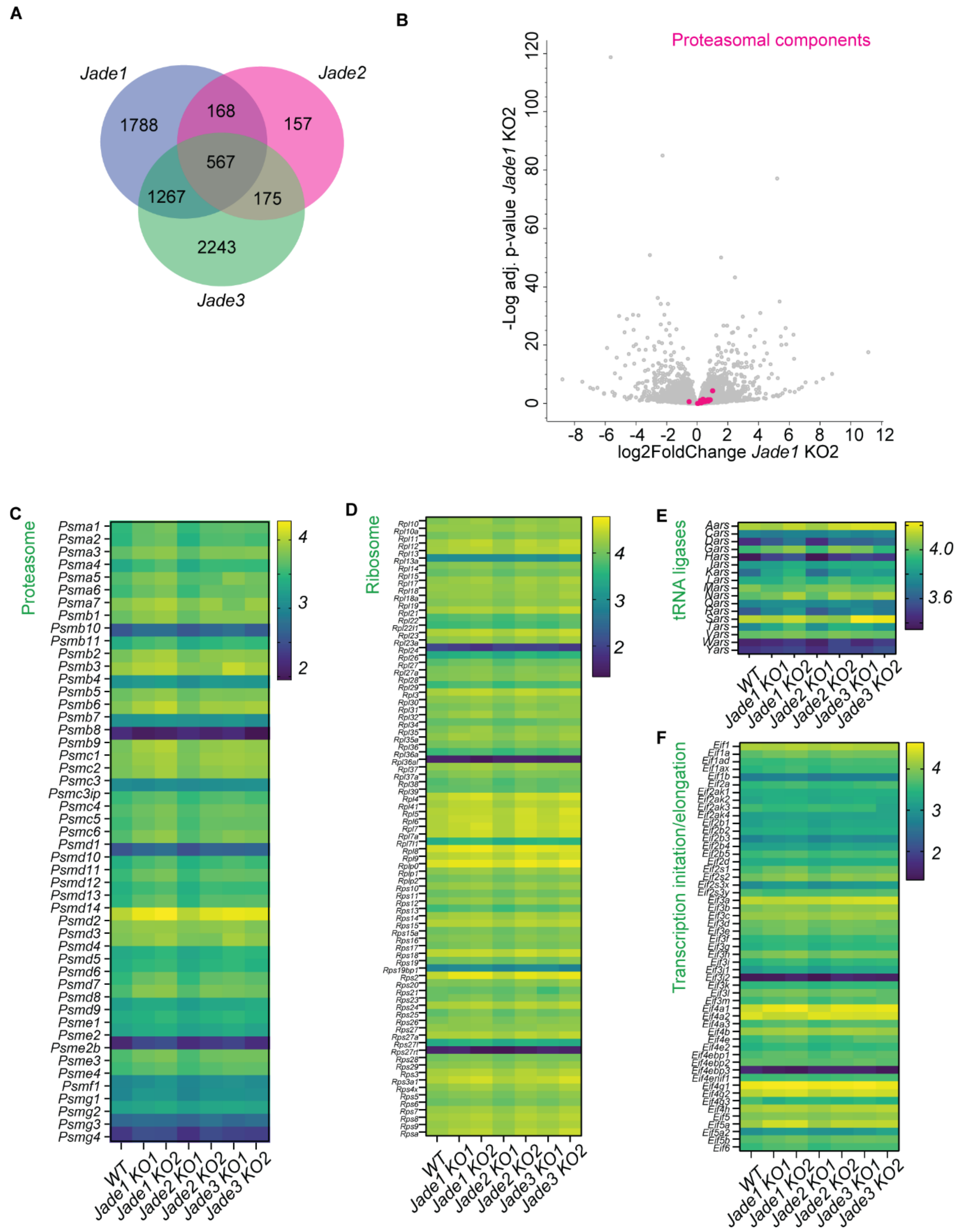


**Supplemental figure S5: Transcriptomic analysis reveals no major regulation of the proteasome. (A)** Venn diagram depicting the overlap between significantly regulated genes across the *Jade*-deficient cell lines. The *Jade1* section includes all proteins that are significantly regulated (p-value < 0.05) in either one or both of the *Jade1*-deficient cell lines. The *Jade2* and *Jade3* sections generated in the same way. **(B)** Scatter plot with the log2FoldChange in gene expression of the *Jade1* KO2 mutant cells vs the wildtype cells on the x-axis and the statistical significance (-log adj. p-value) on the y-axis. Genes of proteasomal components are highlighted in magenta. **(C)** Heatmap of the genes highlighted in (H) based on logarithmic normalized counts for all seven cell lines to visualize that the proteasomal components are not upregulated on a transcriptional level across Jade-deficient cell lines. **(D-F)** Heatmap of ribosomal genes **(D)**, genes encoding for tRNA-ligases **(E)**, and genes encoding for transcription initiation and elongation factors **(F)**. Logarithmic normalized counts for all seven cell lines are shown.

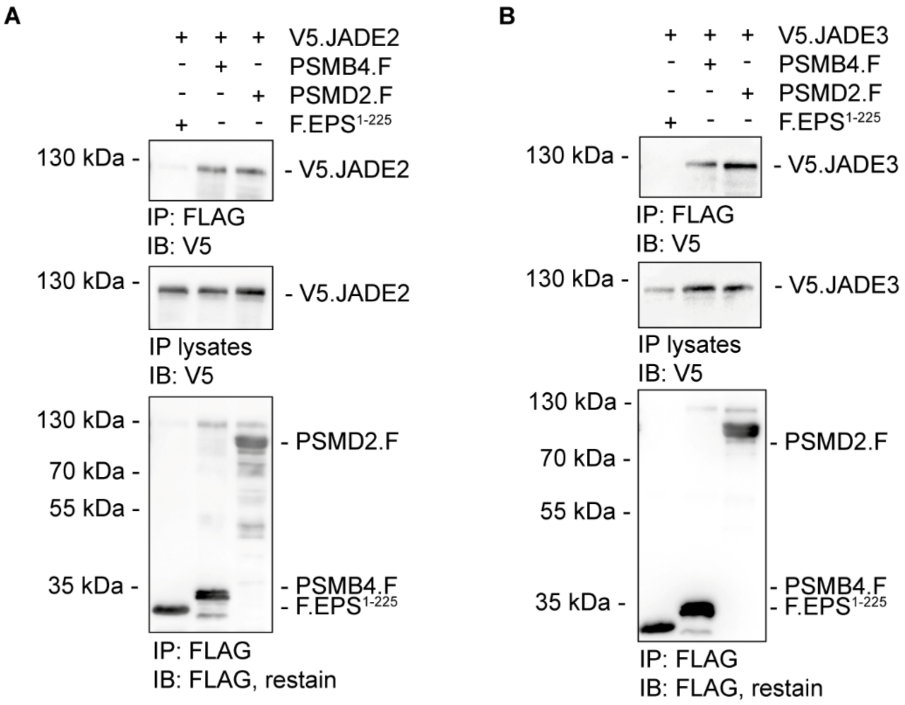


**Supplemental figure S6: Jade2 and Jade3 co-precipitate with the proteasome core components PSMB4 and PSMD2**. **(A/B)** HEK293T cells were transiently transfected with **(A)** V5-tagged JADE2 (V5.JADE2) or **(B)** V5-tagged JADE3 (V5.JADE3), and either FLAG-tagged PSMB4 (PSMB4.F), PSMD2 (PSMD2.F), or a control protein (F.EPS$^{1-225}$). Immunoprecipitates with an anti-FLAG antibody were analyzed by immunoblotting with antibodies (anti-V5; anti-FLAG) as indicated. V5.JADE2 and V5.JADE3 both coimmunoprecipitate with PSMB4.F and PSMD2.F but not with F.EPS$^{1-225}$. Blots are representative for at least three independent experiments.

**Supplemental tables**

**Supplemental table S1: Primer sequences.**

| Name | Sequence 5‘ -> 3‘ | Purpose |
|---|---|---|
| **Jade1_KO1_sgRNA** | ATGGTGTACTCGTCCAGCTC-CGG | sgRNA |
| **Jade1_KO2_sgRNA** | CGACATAGCCAACTGCCCTC-TGG | sgRNA |
| **Jade2_KO1_sgRNA** | GACGTGTGCCGTTCCCCTGA-AGG | sgRNA |
| **Jade2_KO2_sgRNA** | CACTTGGTCCCACTTCTAGT-GGG | sgRNA |
| **Jade3_KO1_sgRNA** | GAACCATGCTATTGAAACAG-TGG | sgRNA |
| **Jade3_Ko2_sgRNA** | GGCATTCTCAAGATTCCAGA-AGG | sgRNA |
| **Jade1_exon6_fp** | AGGATGCTGCGGAAGTTGTT | PCR/sequencing |
| **Jade1_exon6_rp** | ACCCTTTGCTCGGGTTTGTT | PCR |
| **Jade1_exon7_fp** | ACAGCAGGCCATTACTCCAGTT | PCR/sequencing |
| **Jade1_exon7_rp** | CTACTGCTGCCTGTGAGTCACT | PCR |
| **Jade2_exon6_fp** | GCTGAAGCCCAGATCACCAT | PCR/sequencing |
| **Jade2_exon6_rp** | ACTACCCGAAGGACACTTGC | PCR |
| **Jade2_exon7_fp** | TCTCTGTGCCACCCAGTGTA | PCR/sequencing |
| **Jade2_exon7_rp** | GGGAACCCAAGAGCTCCTTCTT | PCR |
| **Jade3_exon5_fp** | AGGGTTTTTATCAGGAGATGGACA | PCR/sequencing |
| **Jade3_exon5_rp** | AGCAGCTTAGAGATCATAAGAAGT | PCR |
| **Jade3_exon6_fp** | CAGATGATGCTACAAGCTTGGTTT | PCR/sequencing |
| **Jade3_exon6_rp** | GAAGACTGGAGGTAGCACAGA | PCR |
| **Jade1_qPCR_fp** | ACCCGCAGCGGAACCAAGTG | qPCR |
| **Jade1_qPCR_rp** | CCGGTGGGCCTCCTCTCGAT | qPCR |
| **Jade2_qPCR_fp** | CCTGTGGTGAGGCTCCTCCC | qPCR |
| **Jade2_qPCR_rp** | CTCTAACGTTAGCTCATCCA | qPCR |
| **Jade3_qPCR_fp** | CAGTGTGCGGCGAGGAA | qPCR |
| **Jade3_qPCR_rp** | ATGCTGGATTCGTCACTCCC | qPCR |
| **Hprt1_fp** | GCTGACCTGCTGGATTACAT | qPCR |
| **Hprt1_rp** | TTGGGGCTGTACTGCTTAAC | qPCR |

**Supplemental table S2: Unique target peptides for PRM assay.**

| Mass [m/z] | Charge state | peptide | Protein |
|---|---|---|---|
| **525.2667** | 2 | VLEEFEQR | Jade1 |
| **566.2982** | 2 | TILAENDEVK | Jade1 |
| **857.1147** | 3 | QKLQQLEDEFYTFVNLLDVAR | Jade1 |
| **723.7175** | 3 | ALRLPEEVVDFLYQYWK | Jade1 |
| **692.859** | 2 | VQEQIFTQYTK | Jade1 |
| **671.7939** | 2 | SLC[CAM]]QEHSDGGPR | Jade2 |
| **559.2766** | 2 | IPEGSWLC[CAM]R | Jade3 |
